# Quantitative Mass Spectrometry Imaging Reveals Mutation Status-independent Lack of Imatinib Penetration into Liver Metastases of Gastrointestinal Stromal Tumors

**DOI:** 10.1101/562553

**Authors:** Denis Abu Sammour, Christian Marsching, Alexander Geisel, Katrin Erich, Sandra Schulz, Carina Ramallo Guevara, Jan-Hinrich Rabe, Alexander Marx, Peter Findeisen, Peter Hohenberger, Carsten Hopf

## Abstract

Mass spectrometry imaging (MSI) is an enabling technology for label-free drug disposition studies at high spatial resolution in life science- and pharmaceutical research. We present the first extensive clinical matrix-assisted laser desorption/ionization (MALDI) quantitative mass spectrometry imaging (qMSI) study of drug uptake and distribution in clinical specimen, analyzing 56 specimens of tumor and corresponding non-tumor tissues from 28 imatinib-treated patients with biopsy-proven gastrointestinal stromal tumors (GIST). For validation, we compared MALDI-TOF-qMSI with conventional UPLC-ESI-QTOF-MS-based quantification from tissue extracts and with ultra-high resolution MALDI-FTICR-qMSI. We introduced a novel generalized nonlinear calibration model of drug quantities based on focused computational evaluation of drug-containing areas that enabled better data fitting and assessment of the inherent method nonlinearities. Imatinib tissue spatial maps revealed striking inefficiency in drug penetration into GIST liver metastases even though the corresponding healthy liver tissues in the vicinity showed abundant imatinib levels beyond the limit of quantification (LOQ), thus providing evidence for secondary drug resistance independent of mutation status. Taken together, these findings underline the important application of MALDI-qMSI for studying the spatial distribution of molecularly targeted therapeutics in oncology.

## Introduction

Matrix-assisted laser desorption/ionization (MALDI) mass spectrometry imaging (MSI) has emerged as a powerful label-free technology for studying spatial distributions of analytes in tissues. Owing to its high chemical specificity and ability to track hundreds of analytes simultaneously, MALDI-MSI has gained traction in various areas of biomolecular research such as quantitative profiling of metabolites, lipids, peptides, proteins and drugs (1–4). In the latter case, MALDI-MSI has found its way into pharmaceutical research and development, where disposition of drugs and their carriers can be effectively monitored alongside their pharmacodynamics and toxic effects (4–8).

Traditionally, quantitative assessment of drug distribution in patient tissues employs bioanalytical techniques such as UPLC-ESI-MS, which assume homogenous distribution of analytes and require tissue homogenization resulting in a complete loss of spatial context (5). In contrast, MALDI-MSI provides spatial information, and much effort has recently been devoted to establish quantitative MSI (qMSI) techniques (1–4, 9–12). Linear calibration based on tissue mimetic models or compound dilution series spotted onto tissue, as well as signal normalization against stable isotope-labeled internal standards (IS) (11) and the calculation of tissue extinction coefficients (TEC) (10) were introduced to compensate for the inherently high technical variability of MALDI-MSI (3, 4, 12). Furthermore, advanced rational testing criteria for evaluating linearity of response, variability, reproducibility and limits of detection of qMSI have been suggested (13). Despite these technical advances, MALDI-qMSI has not yet been widely adapted to clinical pharmacology.

In clinical oncology, the ability to monitor drug penetration into tumors constitutes a key medical need (14). However, tumor heterogeneity and the poorly understood spatial organization of the tumor microenvironment present major challenges for drug uptake and, hence, effective cancer treatment (15, 16). This challenge has prompted qMSI studies of drug disposition and of pharmacological/toxic effects in tumor tissues and their surroundings in mice (17, 18). Most mouse studies and pioneering qualitative MSI studies of the tyrosine kinase inhibitor (TKI) erlotinib in patient tissue report high degrees of variability and intratumor heterogeneity as well as highly heterogeneous drug distribution (19–21). However, clinical qMSI proof-of-concept studies of drug distribution are still lacking.

Here, we present the first systematic MALDI-qMSI drug disposition study of a TKI, imatinib, in 56 resection specimens of tumor and surrounding non-tumor tissues from 28 patients with biopsy-proven gastrointestinal stromal tumor (GIST). Imatinib is the standard first line treatment for GIST, effectively stopping autophosphorylation and tumor proliferation particularly in exon 11 mutated GIST (22–24). However, tumors are prone to imatinib resistance, which is mainly attributed to secondary somatic mutations in the KIT or the platelet-derived growth factor receptor alpha (PDGFRA) tyrosine kinases (25, 26). In this study, we introduce generalized nonlinear regression as a superior calibration method based on imatinib-containing pixels, report MALDI-qMSI of three full technical replicates, and compare these results [fast time of flight MALDI-TOF-qMSI and ultra-high-resolution Fourier-Transform Ion Cyclotron Resonance (FTICR-qMSI)] with conventional UPLC-ESI-QTOF-MS quantification. Taken together, our data suggests that MALDI-qMSI compares well with UPLC-ESI-QTOF-MS quantification, and that spatially resolved MS has utility in clinical pharmacology. We can demonstrate that independent of mutation status of the tumor, imatinib failed to penetrate or to be retained in tumor tissue in all GIST liver metastasis cases tested.

## Methods (see also Supporting Information)

### Tissue Samples

Human GIST and corresponding non-tumor control tissues (total of 56 specimens) had been surgically removed from 28 patients (**Supporting Information; Table 1**) with approval by the Medical Ethics Committee II of the Medical Faculty Mannheim of Heidelberg University (2012-289N-MA; 2015-868R-MA; 2017-806R-MA). All patients had been treated with 400-800 mg imatinib orally once daily including the day prior to surgery, hence, tumor and healthy tissue samples were analyzed at a steady state drug status before surgery. After removal, tumor tissue was cut into small pieces, snap-frozen and stored at −80 °C in the biobank of the Department of Surgery at University Hospital Mannheim. For all samples, histology, mitotic activity, regression index and CD117 (=*KIT*), CD34 as well as DOG1 immunohistochemistry (IHC) were assessed. Exons 9, 11, 13, 14 and 17 of the *KIT* gene as well as exons 12 and 18 of the *PDGFRA* gene were analyzed for mutations after tumor tissue microdissection. Porcine liver tissue was used to establish drug calibration curves on test slides. These were supplied by the local slaughterhouse and immediately snap-frozen (−80°).

### Experimental Setup

Each sample was sectioned in six steps at –20 °C in a CM 1950 cryostat (Leica Biosystems GmbH, Nussloch, Germany) in triplicate (see **Supporting Information;** Figure 1a). After trimming, the first 8-µm tissue section of each sample was thaw-mounted onto a gold target for MALDI-TOF-qMSI. Two consecutive slices were then put onto Starfrost adhesive microscope slides (R. Langenbrinck GmbH, Emmendingen, Germany) for standard histological analysis (one stained with hematoxylin & eosin (H&E) and one backup slide) and scanned in an Aperio CS2 (Leica Biosystems). Afterwards, four consecutive sections were collected for quantitative drug determination by UPLC-ESI-QTOF-MS. Finally, some exemplary sections were mounted on conductive indium tin oxide (ITO) slides for additional MALDI-qMSI with a high mass-resolving FTICR-MS (n = 3). For MALDI-qMSI, each ITO slide featured duplicate spots of an imatinib dilution series (25, 12.5, 6.25, 3.125, 1.56, 0.78 pmol and a blank control) spotted onto a porcine liver section. For a schematic overview of the arrangement of tissue sections on slides, see **Supporting Information;** Figure 1b). To achieve comparable conditions after preparation, all slides were stored in a desiccator at room temperature for one hour.

**Figure 1.**
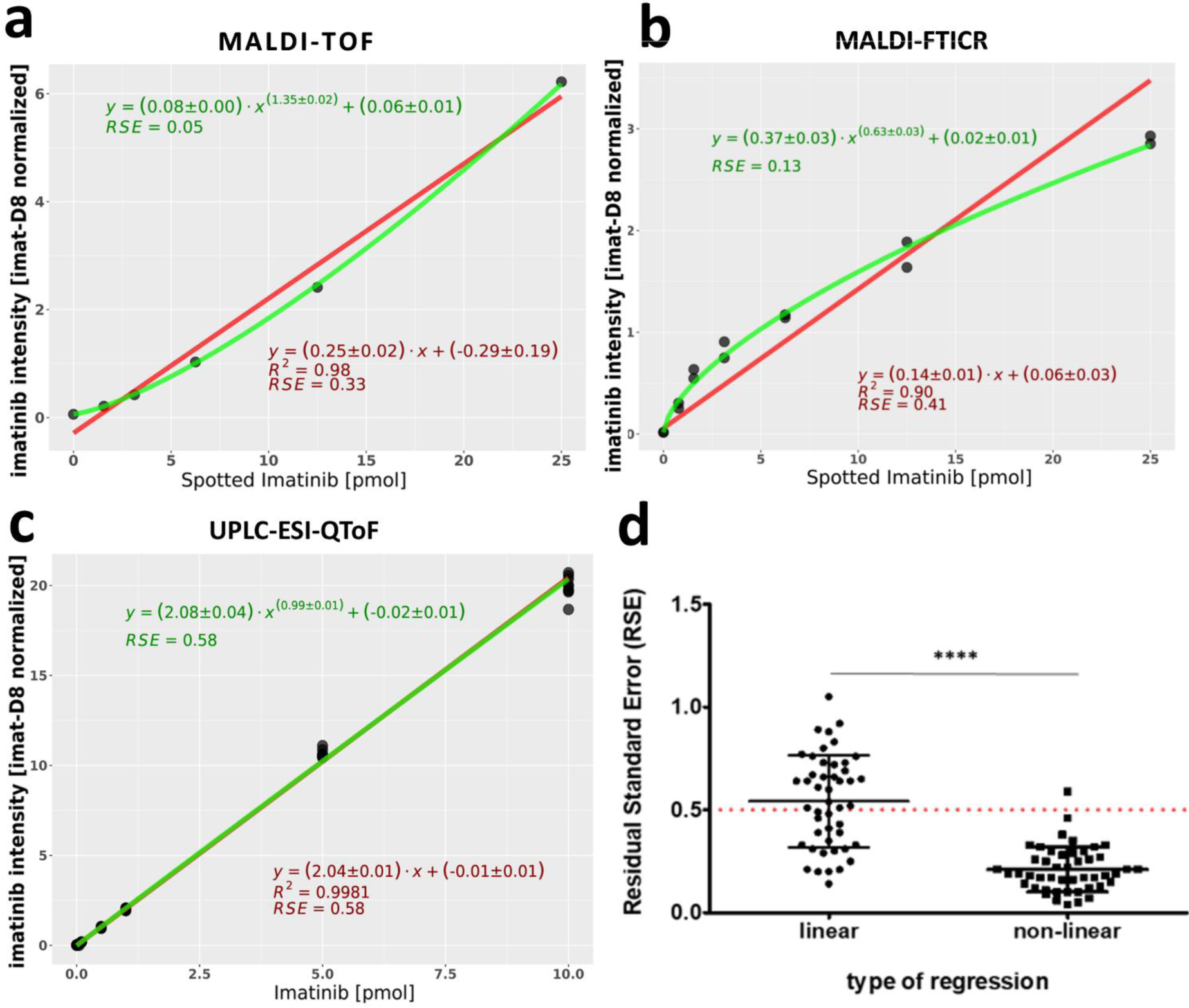
Generalized nonlinear regression model for calibration in qMSI. Comparison of calibration curves for a sample dataset by linear (red) and generalized nonlinear regression (green) for **a)** MALDI-TOF-qMSI, **b)** MALDI-FTICR-qMSI and **c)** UPLC-ESI-QTOF-MS. Grey circles represent the mean intensity of the imatinib-containing pixel within a dilution series area. **d)** RSE for linear and generalized nonlinear calibration curves of MALDI-TOF-qMSI.

### Quantitative MALDI-TOF- and -FTICR-qMSI Imaging

For normalization, nine layers of deuterated imatinib (imatinib-D8) were deposited on the slides using a SunCollect sprayer (SunChrom GmbH, Friedrichsdorf, Germany): flow rate 15 µL/min, spray head distance 45 mm and speed 300 mm/min. Slides were stored in a desiccator at room temperature for one hour to allow drying of the IS. For positive-ion mode measurements, five layers (flow rates of 10, 15, 3 × 20 μL/min) of 2,5-Dihydroxybenzoic acid (DHB) matrix (60 mg/mL in 50% acetonitrile, 0.5% Trifluoroacetic acid) were subsequently sprayed onto the tissue at 300 mm/min as described previously (27).

MALDI-TOF-qMSI data acquisition was performed on an ultrafleXtreme MS using flexControl 3.4 (Bruker Daltonics GmbH; for instrument parameters, see **Supporting Information; section 1.3**). High-resolving-power MSI (*m/z* 150–3000; R: 260,000 at *m/z* 300) was recorded in positive-ion mode on a solariX 7T XR FTICR mass spectrometer (Bruker Daltonics). The raster width was 200 μm for calibration curves for quantification and 50 μm for all human samples. The instrument was calibrated externally using quadratic mass calibration with peptide calibration standard II including dasatinib (*m/z* 488.21). For imatinib calibration curves, equal square-sized measurement areas containing identical pixel numbers were defined over each calibration spot by an in-house MATLAB tool.

### DATA IMPORT, CONVERSION AND PRE-PROCESSING

A total of 48 MALDI-TOF- and three MALDI-FTICR-qMSI-datasets were acquired and exported into imzML format (28) using flexImaging 4.1 (Bruker Daltonics) with a binning rate of 120,000. The imzML datasets were subsequently imported into R 3.3.1 (R Foundation for Statistical Computing, Vienna, Austria)(29) using *MALDIquantForeign* and processed using *MALDIquant* packages (30). Mass spectra were normalized to the maximum peak intensity over the mass range *m/z* 502.32 ± 100 *ppm* (MALDI-TOF-qMSI) and ± 10 *ppm* (MALDI-FTICR-qMSI) of the sprayed IS imatinib-D8. Neither baseline correction nor further pre-processing of spectra were performed. All subsequent analysis and visualization (mainly with *ggplot2* and *fmsb* packages) were performed using R.

### REGRESSION MODELS AND QUANTIFICATION

Linear regression analysis for calibration of the spotted imatinib dilution series was performed as described (9, 10). Nonlinear regression was performed by fitting a power function as a calibration curve in the form of *y* = *a x^b^* + *c* model (for details see **Supporting Information; section 1.2**). For MALDI-qMSI of imatinib in tissue, the mean intensity of signal-bearing pixels, i.e. imatinib signal with signal to noise ratio (S/N) ≥ 3, of each tissue section was plugged into the fitted nonlinear regression model. The limits of detection (LOD) and quantification (LOQ) in this study were computed as mean of all individually fitted calibration curves (48 for MALDI-TOF-qMSI and 3 for MALDI-FTICR-qMSI; see **Supporting Information**). To assess imatinib ion suppression within different tissues, TEC (10) was calculated as the ratio of the mean intensity of the deuterated IS (m/z 502.32 ± 100 *ppm*) within each tissue section to its mean intensity in the IS and matrix control area. The residual standard error of the calibration curve (RSE) was used for quality of fit comparison, as R2 is not defined for nonlinear fits (31).

## Results

### Generalized nonlinear regression for calibration in MALDI-ToF- and -FTICR-qMSI

For clinical MALDI-qMSI proof-of-concept, we systematically examined GIST tumor- and corresponding non-tumor samples from 28 patients who received the TKI imatinib as first line treatment but presented with refractory disease. Some patients had developed metastatic lesions, mostly in the liver (**Supporting Information; Table 1**). All patients received imatinib prior to surgery according to their treatment plan, thus enabling investigation of the drug’s distribution in resection specimens. Due to the limited number of available tissue samples and since all measurements were conducted in triplicates, only samples from 18 and 5 patients were available for UPLC-ESI-QTOF-MS and MALDI-FTICR-qMSI measurements, respectively. All cryopreserved tissues were cut into several adjacent sections for MALDI-qMSI, for H&E histology, IHC, and for compound extraction and subsequent UPLC-ESI-QTOF-MS quantification (n=3 each; **Supporting Information;** Figure 1a **and** 2). All slides for MALDI-qMSI featured an imatinib dilution series spotted onto porcine liver tissue and a spray-coating of deuterated imatinib-D8 for lock-mass calibration and normalization, respectively (**Supporting Information;** Figure 1b).

**Figure 2.**
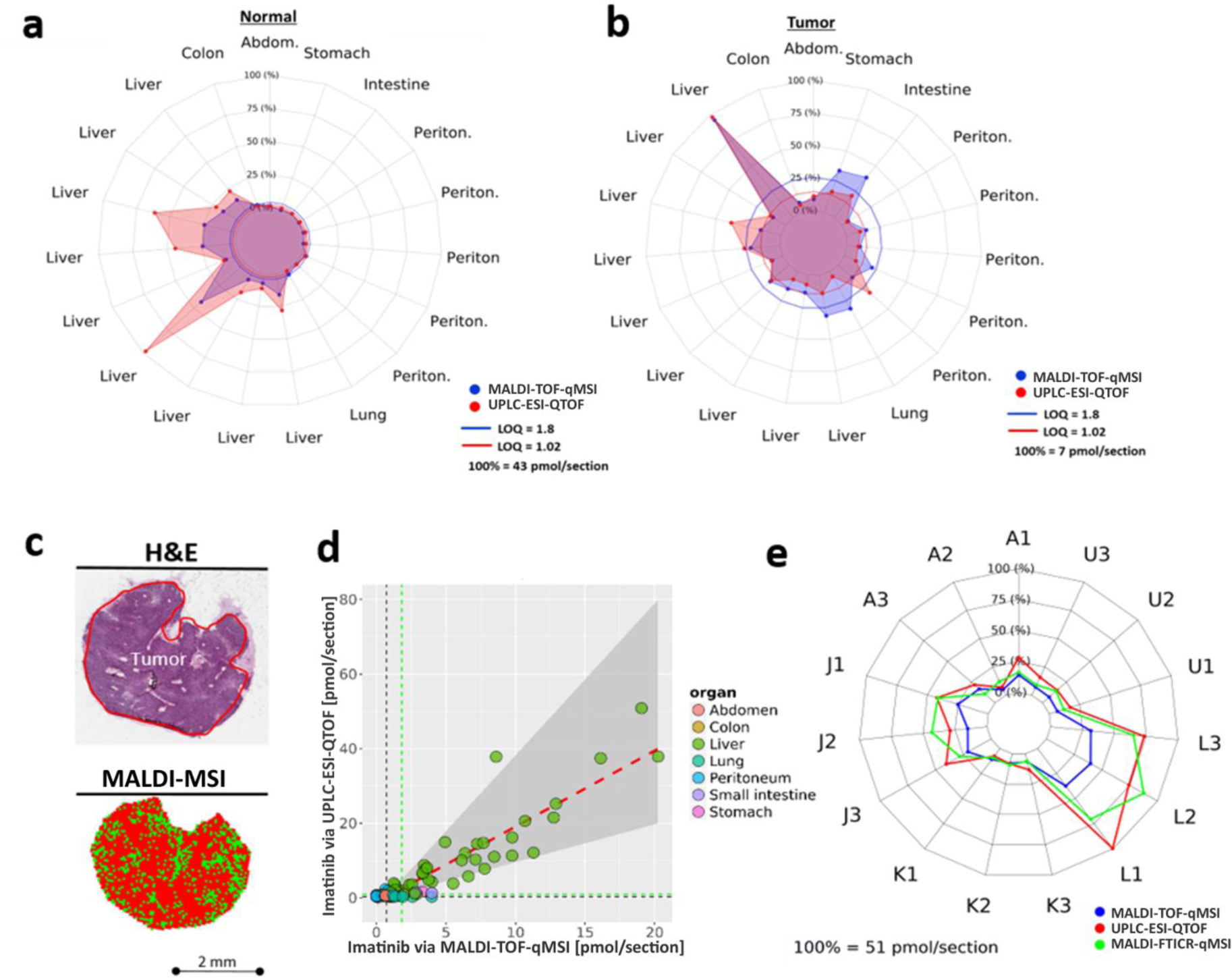
Verification of MALDI-TOF-qMSI for imatinib quantification in GIST. Comparison of MALDI-TOF-qMSI and UPLC-ESI-QTOF-MS quantification for **a)** normal and **b)** tumor tissue. **c)** A liver metastasis (sample J) illustrating the drug-containing pixels (bottom) and corresponding H&E-stained image (top). **d)** Correlation of quantification between MALDI-TOF-qMSI and UPLC-ESI-QTOF-MS. **e)** Comparison of MALDI-TOF-qMSI, UPLC-ESI-QTOF-MS and MALDI-FTICR-qMSI quantification for five (x3) unaffected liver samples.

For drug quantification, a calibration curve has to be fitted to this dilution series (4), and users typically manually encircle spots that are then used as regions of interest in computation of the calibration curve. In contrast, we computationally defined drug-bearing areas, including only pixels where the drug was detected with S/N ≥ 3 (**Supporting Information;** Figure 3). The lower the drug concentration spotted, the more the outlines of drug-spotted areas diverge from the expected circular shape (**Supporting Information;** Figure 3a). This can lead to signal dilution and underestimation of signal intensity when the mean signal intensity includes pixels that do not carry drug signal (**Supporting Information;** Figure 3b). This effect is even more drastic in heterogeneous tumor tissue, where the drug can only be observed in a few pixels.

**Figure 3.**
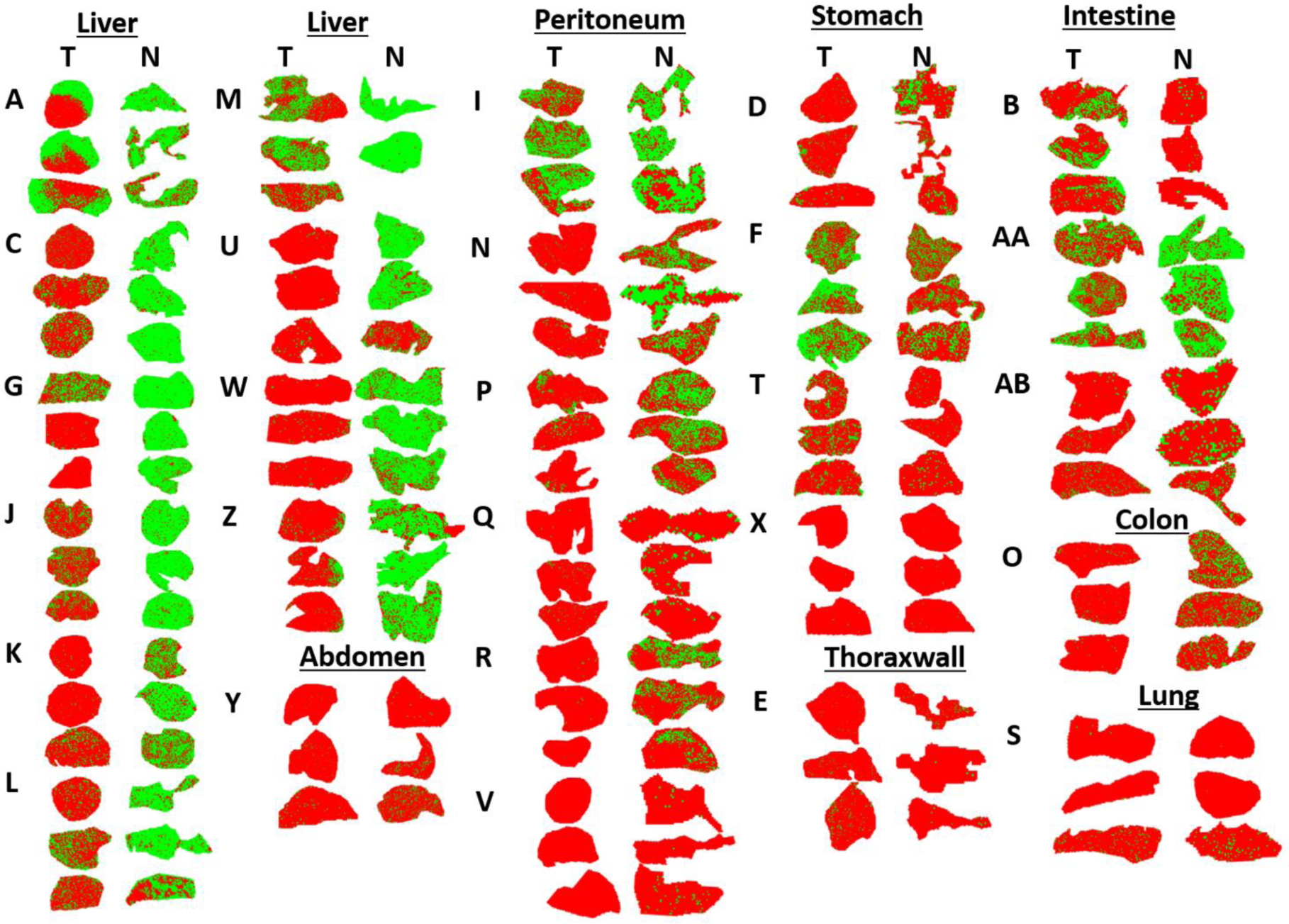
Imatinib distribution in all tumor (T) and normal (N) non-tumor samples from GIST patients. Green and red pixels indicate imatinib signal presence (S/N ≥ 3) and absence, respectively. Three cryosections were prepared per tissue sample/patient, and patients are coded by single or double letters. Tissues identified as stomach, colon or intestine are primary tumors. All others were metastases.

### Quantitative analysis of imatinib content in GIST tumor specimens by UPLC-ESI-QTOF-MS and MALDI-qMSI

More importantly, most drug dilution series measured either with MALDI-TOF-qMSI or MALDI-FTICR-qMSI (**Supporting Information;** Figure 4) deviated from linearity. Similar nonlinear behavior of calibration curves has been reported by others (11). We therefore wondered if nonlinear regression fitting might be a better option for calibration than a linear regression fit. As illustrated for exemplary dilution series, the generalized nonlinear calibration provided a much better fit indicated by the 6- and 3-fold better RSE for MALDI-TOF-qMSI and MALDI-FTICR-qMSI (Figure 1a,b), respectively. Figure 1d indicates a significant decrease in RSE for all MALDI-TOF-qMSI calibration curves (p<0.0001; n=48) for the nonlinear calibration when compared to the linear one. The linear and nonlinear fits were nearly identical for UPLC-ESI-QTOF-MS calibration (Figure 1c) indicating a true linear response of the system

**Figure 4.**
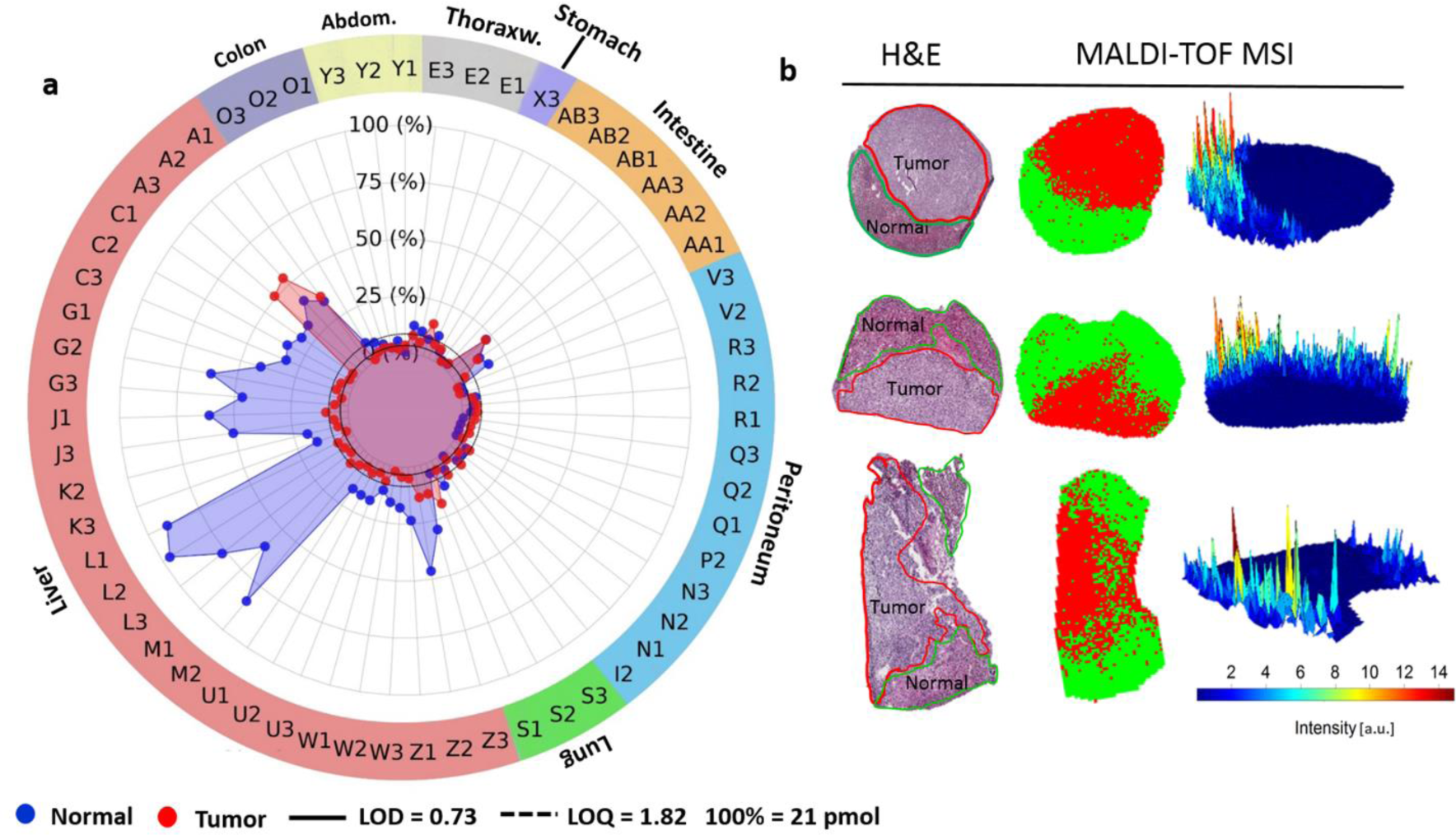
Liver metastases of GIST display limited imatinib content independent of mutation status. **a)** MALDI-TOF-qMSI-Quantified imatinib in GIST samples cohort comparing “Tumor” (red) and corresponding “Normal” (blue) tissues. (LOD = 0.73 pmol/section; LOQ = 1.82 pmol/section.) **b)** Three sample A replicates containing both “normal” and “tumor” tissue based on histopathological re-examination Illustrating imatinib’s absence from metastatic GIST.

Next we systematically compared the average imatinib content in both normal and tumor tissue sections (in pmol/section) for all three full replicates using gold-standard UPLC-ESI-QTOF-MS and MALDI-TOF-qMSI (Figure 2a,b). Also for this comparison, the mean imatinib signal intensity in MALDI-qMSI datasets was calculated solely from imatinib-containing pixels (S/N ≥ 3 for *m/z* 494.26; Figure 2c). Most tissue sections with imatinib levels above LOQ for both MS methods corresponded to normal liver tissue. In these cases, imatinib quantified by MALDI-TOF-qMSI displayed acceptable correlation with results from UPLC-ESI-QTOF-MS quantification. However, MALDI-TOF-qMSI tended to underestimate imatinib. Nevertheless, 78% of all samples, in which imatinib could be quantified (in Figure 2d; upwards and rightwards of the green-coloured dashed LOQ lines [MALDI-TOF-qMSI; 1.8 pmol/section, UPLC-ESI-QTOF-MS; 1.0 pmol/section]), were inside a window of a 2-fold difference (grey area in Figure 2d). MALDI-FTICR-qMSI matched the UPLC-ESI-QTOF-MS results even more closely (Figure 2e).

### Limited imatinib uptake in metastatic GIST independent of mutation status

To be effective, cancer-targeting drugs must adequately penetrate into tissue. Hence, we sought to investigate imatinib’s tumor penetration capability in human GIST samples. Figure 3 illustrates an overview of the measured tissue sections for all 28 patient samples, done in triplicates, with green and red pixels indicating detected (S/N ≥ 3) and undetectable drug signal, respectively (also see **Supporting Information; Table 1** for clinical and histopathological information and **Supporting Information;** Figure 2 for H&E-stained sections). Follow up histopathological examination (**Supporting Information;** Figure 2) suggested that 26 of the tumor tissue replicates examined contained only slight traces of regressive tumor areas with fibrosis and necrotic tissue throughout. They were, therefore, omitted from further analysis.

Systematic quantification of imatinib (pmol/section) in tumor tissue sections and their corresponding non-tumor (normal) control tissues by MALDI-TOF-qMSI revealed that despite continuous administration of the prescribed dosage of imatinib until surgery (see **Supporting Information; Table 1**), tissue drug levels were typically below LOQ (Figure 4a; but see sample AA as an exception).

In most tumor sections (48 of 60 sections, 80%, in MALDI-TOF-qMSI and 31 of 44, 70%, in UPLC-ESI-QTOF-MS), regardless of its type (i.e. primary tumor or metastasis) and mutation status, imatinib content was below LOQ. In comparison, for the corresponding normal tissue sections only 40 of 83 sections (48%) and 23 of 54 sections (43%) contained imatinib amounts below LOQ for MALDI-TOF-qMSI and UPLC-ESI-QTOF-MS, respectively. Imatinib levels were higher in normal liver tissues compared to the others.

Surprisingly though, the orally administered imatinib had limited uptake or retention in metastatic GIST in liver tissue, leading to amounts below LOQ despite the high abundance of the drug (well above LOQ) within surrounding normal liver tissue. *Sample A* appears to be a notable exception, as comparable amounts of imatinib were present in both normal and tumor tissue (also observed with UPLC-ESI-QTOF-MS in **Supporting Information; Figure 7**). However, upon histopathological re-investigation, all three replicates of that sample were found to contain both normal hepatic and metastatic GIST tissues (Figure 4b). As revealed by spatially-resolved-qMSI, also this sample contained imatinib only in its non-tumor part (Figure 4b). Apparently, the drug was unable to penetrate into the metastatic tumor despite high concentrations within the surrounding tissue. To rule out the possibility that observed differences in imatinib signal were simply a consequence of different desorption/ionization characteristics of the respective tissues, we calculated TEC for all normal tissue-tumor tissue pairs (**Supporting Information; Figure 5**): Importantly, for liver, stomach and lung no significant differences between the drug’s ionization for normal and corresponding tumor tissue were observed. Additionally, no other interfering analytes were detected within 100 *ppm* of imatinib as verified by high mass resolution MALDI-FTICR-MSI as shown in **Supporting Information; Figure 6**. Hence, MALDI-TOF-qMSI results reflect real differences in drug distribution.

## Discussion

For successful therapy, anticancer drugs must penetrate into tumor tissue and reach their target in cancer cells at sufficient quantities. In this study, we aimed to improve computational calibration in MALDI-TOF-qMSI, to provide proof-of-concept for clinical utility of this technique by rigorously comparing it with conventional UPLC-ESI-QTOF-MS-based quantification and to cast light onto the effectiveness of uptake (or lack of export/intra-tissue metabolism) of the therapeutic TKI, imatinib, into GIST tissue obtained from 28 patients.

In contrast to UPLC-ESI-QTOF-MS measurements that utilized extracts of entire tissue slices, MALDI-TOF- and –FTICR-qMSI add the crucial benefit of delivering spatial drug distribution maps, which - with histopathological tissue annotation - enables co-localization analysis of drugs within different tissue morphologies. Loss of spatial information puts constraints on UPLC-ESI-QTOF-MS measurements of “tumor tissue”, which are always skewed by tissue heterogeneity, i.e. the presence of unknown fractions of non-tumor tissues in tumor samples. S*ample A*, for which UPLC-ESI-QTOF-MS showed similar levels of imatinib in normal and tumor tissue, serves as point in case (**Supporting Information; Figure 7**), which was explained by H&E annotation and MALDI-TOF-qMSI images (Figure 4). Moreover, even though MALDI-TOF- and - FTICR-qMSI has been traditionally referenced to UPLC-ESI-QTOF-MS as gold standard for absolute quantification, such validation by UPLC-ESI-QTOF-MS needs to be considered carefully, since MALDI-qMSI reports pixel-wise quantification of few nm thin surface layer (32), while the latter quantifies the drug within a defined tissue volume.

For proper quantification, the systematic nonlinearity observed in all 48 and 3 dilution sets (**Supporting Information;** Figure 4) of MALDI-TOF- and -FTICR qMSI, respectively, is particularly relevant. The same behavior has already been reported by Pirman and co-workers (11), and was attributed to the matrix-to-analyte ratio. Moreover, the complex MALDI process, non-uniform tissue-ion suppression, and interferences from matrix background signals are all factors that could contribute to nonlinear responses (33). Even though it is common for MALDI-TOF- and –FTICR-qMSI studies to use linear calibration for drug dilution series, this linearity cannot always be guaranteed for rare heterogeneous tissue samples even with optimized matrix-to-analyte ratios (see (10)). Furthermore, especially in a clinical screening setting where samples are obtained from different individuals and contain substantially different amounts of target analyte, it is unlikely that any method can guarantee optimal matrix-to-analyte ratios in all scenarios and for all samples. Therefore, a linear behavior of calibration curves for all samples cannot be guaranteed. Hence, nonlinear calibration seems to be the method of choice. On another cautionary note, we suggests that for a reliable reporting of a drug’s uptake and distribution, the level of the drug should always be reported relative to its LOQ; drawing conclusions based on low intensity MS ion images of the drug can be potentially misleading and unreliable.

For GIST metastases in liver, the discrepancy between imatinib amounts in normal hepatic versus metastatic tissue is striking (e.g. replicates of *sample A* (Figure 4b)). Further work is required to elucidate if lack of imatinib in liver metastases reflects effective export/metabolism or inefficient uptake. For instance, higher expressions of P-glycoprotein and multiresistance protein 1 (MRP1), which are involved in active efflux of a wide spectrum of drugs including imatinib, have been reported in gastric and nongastric GISTs, respectively (34). Other, rather passive, reasons for such drug-resistance that is independent of the KIT and PDGFRA mutation types, a poorly organized vasculature, increased interstitial fluid caused by the lack of functional lymphatics and/or inflammation and an abnormal structures of the extracellular matrix (14, 15). Additionally, it is well-known that tumor vascularity decreases in GIST metastases after imatinib treatment, which may lay the basis for subsequent tumor progression due to inadequate local drug concentrations (35, 36). The observed apparent lack of drug also confirms the clinical practice for GIST patients with liver metastases that the best treatment option is a multidisciplinary one with continued TKI drug therapy and possibly surgical intervention (37). This result based on a small cohort of patient-derived tissues is in line with results from several MALDI-qMSI mouse studies on various chemotherapeutic drug agents in different cell line-based and patient-derived xenograft models (18, 19, 21, 38–40).

## Conclusion

While providing additional spatial information on imatinib distribution, MALDI-qMSI based on generalized nonlinear calibration and focused computational analysis of drug-bearing pixels provided quantitative measures of imatinib that deviated from reference analysis by UPLC-ESI-QTOF-MS by less than two-fold in 78% of cases and provided a generalized method for modeling analytes dilu-tion series as well as assessing the degree of the apparent nonlinearity of the system. While further refinement of the method and testing in larger patient cohorts will be crucial, our data provides proof-of-concept for clinical utility of MALDI-qMSI. Furthermore, spatial mapping of imatinib distribution within GIST patient tissues revealed striking inefficiency in its penetration into liver metastases irrespec-tive of their mutation status. Our study therefore strongly suggests that also mechanisms other than driver mutations in receptor tyrosine kinases such as alterations in drug up-take, efflux or intratumor metabolism may play an im-portant role in imatinib resistance in GIST. This finding underlines the important application of MALDI-qMSI for studying the spatial distribution of molecularly targeted therapeutics in oncology.

## Acknowledgements

This work was supported by the European Union’s Seventh Framework Programme (FP7/2007–2013) under grant agreement no 602306 (MITIGATE; to C.H.). This research project was also funded by the German Federal Ministry of Education and Research (BMBF) within the Framework “Forschungscampus: public-private partnership for Innovations” (M2oBiTE grant 13GW0091B to C.H. and 13GW0091E to A.M.). The authors thank Karin Berner-Leischner for technical assistance with the tissue samples. Eva Wardelmann, Dept. of Pathology, University Hospital Münster was instrumental in analyzing secondary resistance mutations. C.H. thanks the Deutsche Forschungsgemein¬schaft (DFG) for the funding of a SolariX FTICR MS (INST 874/7-1 FUGG).

## Abbreviations

MALDI: Matrix-assisted laser desorption/ionization
MSI: mass spectrometry imaging
UPLC: ultra-performance liquid chromatography
ESI: electro-spray ionization
QTOF: quadruple TOF
MS: mass spectrometry
qMSI: quantitative MSI
TEC: tissue extinction coefficient
TKI: tyrosine kinase inhibitor
GIST: gastro-intestinal stromal tumor
FTICR: Fourier-Transform Ion Cyclotron Resonance
IHC: immunohistochemistry
H&E: hematoxylin & eosin
ITO: indium tin oxide
DHB: 2,5-Dihydroxybenzoic acid
LOD: limit of detection
LOQ: limit of quantification
RSE: residual standard error
S/N: signal-to-noise ratio
MRP1: multiresistance protein 1

## Human Genesw

PDGFRA: platelet-derived growth factor receptor alpha
KIT: KIT proto-oncogene receptor tyrosine kinase

