## Supporting Information (PDF) for "Quantitative Mass Spectrometry Imaging Reveals Mutation Status-independent Lack of Imatinib Penetration into Liver Metastases of Gastrointestinal Stromal Tumors"

#### Table of Contents

**1. Supplemental Methods** ... 2

**1.1.** Chemicals and Reagents ... 2

**1.2.** Calibration Curves ... 2

**1.3.** MALDI-TOF and –FTICR instrument settings ... 5

**1.4.** Tissue Extraction and Imatinib Quantification by UPHPLC-MS Analysis ... 5

**2. Supplemental Tables** ... 7

Supplemental Table 1 ... 7

**3. Supplemental Figures** ... 8

Supplemental Figure 1 ... 8

Supplemental Figure 2 ... 9

Supplemental Figure 3 ... 10

Supplemental Figure 4 ... 11

Supplemental Figure 5 ... 12

Supplemental Figure 6 ... 12

Supplemental Figure 7 ... 13

**4. References** ... 14

### 1. Supplemental Methods

#### 1.1. Chemicals and Reagents

Imatinib (m/z 494.266) and deuterated imatinib-D8 (m/z 502.316, as internal standard; IS) were purchased from LC Laboratories (Woburn, USA) and AlsaChim (Illkirch, France), respectively. Gold-coated slides were from Science Services (Munich, Germany). Indium tin oxide (ITO)-coated glass slides, 2,5-dihyroxybenzoic acid (DHB) and peptide calibration standard II were purchased from Bruker Daltonics (Bremen, Germany). Dasatinib (m/z 488.163; 25 µM) was added to the calibration mix. Trifluoroacetic acid (TFA), xylole and absolute ethanol were from Merck (Darmstadt, Germany); acetone, acetonitrile and methanol from VWR Chemicals (Fontenay-sous-Bois, France); formic acid and Mayer's hematoxylin solution from Sigma-Aldrich (Steinheim, Germany); Eosin G Solution 0.5 from Carl Roth (Karlsruhe, Germany), and DMSO was from Applichem (Darmstadt, Germany). All solvents used were MS grade.

#### 1.2. Calibration Curves

**Linear Model.** In order to quantify the amount of imatinib within the patient tissue samples a calibration curve had to be fit to the imatinib dilution series. In accordance with the current literature[1,2], a linear regression has been attempted to find the linear relation between Imatinib signal intensity $y$ ($a.u.$; normalized to the internal standard) and amount of spotted Imatinib $x$ ($pmol$) such that

$$y = a\,x + \ b \qquad (1)$$

Where $a$ is the slope and $b$ is the $y$-intercept. Let $y_{Limit}$ denote the smallest detectable signal intensity i.e. the limit of detection (LOD), $\bar{y}_{Blank}$ and $\sigma_{Blank}$ are the mean and the standard deviation of the signal intensity in the blank (control) measurement area, respectively. Assuming a normal distribution of the noise signal it can be stated that

$$y_{Limit} = \bar{y}_{Blank} + k\,\sigma_{Blank} \quad (2)$$

Where k is a factor normally equal to $3$ for a confidence level $> 99\%$. In other words, the smallest reliably detectable drug signal is $k = 3$ standard deviations above the mean signal, all measured within the blank (control) measurement area. Using the same rational equation (1) can be re-written to reflect LOD

$$y_{Limit} = a\,x_{Limit} + \bar{y}_{Blank} \quad (3)$$

Rewriting equation (3) as a function of $y_{Limit}$ and plugging equation (2) into it yields

$$x_{Limit} = \frac{k\,\sigma_{Blank}}{a} \quad (4)$$

Where $x_{Limit} = LOD$ for $k = 3$ and $x_{Limit} = LOQ$ for $k = 10$. In other words, the LOD and LOQ for a linear regression model can be expressed (here, in the unit of $pmol$) conveniently by the standard deviation of the signal of interest in the blank and the slope of the fitted model. Moreover, $\sigma_{Blank}$ can also be approximated by the standard error of the $y$-intercept in the fitted linear model.

To limit the effect of heteroscedasticity on the regression model, which is normally observed in MALDI-MSI[3–5], a weighted linear regression has been performed such that each calibration point (representing the mean intensity of the drug signal within the respective dilution area) was down weighted by its pixel-wise variance within that area.

**Generalized nonlinear model.** A nonlinear behavior of the Imatinib dilution series which closely resembles a power-function response was observed in all measured datasets. A similar behavior had been noted in a previous study [6]. Therefore, a nonlinear regression was performed by fitting a power function as a calibration curve in the form of

$$y = a\, x^b + c \tag{5}$$

Where $a$ and $b$ are constants and $c$ was added to represent the superimposed noise error (background signal when the drug signal is absent). Note that as $b \rightarrow 1$, the model's response approaches linearity and the above equation reduces to an equation of a line. To quantify the LOD and LOQ, equation (5) can be re-written as follows

$$y_{Limit} = a\, x_{Limit}^b + \bar{y}_{Blank} \tag{6}$$

Rearranging the above equation a function of $y_{Limit}$ and plugging equation (2) into it yields

$$x_{Limit} = \left(\frac{k\, \sigma_{Blank}}{a}\right)^{1/b} \tag{7}$$

Where $x_{Limit} = LOD$ for $k = 3$ and $x_{Limit} = LOQ$ for $k = 10$. Therefore, similar to the linear model, the LOD and LOQ for the nonlinear fit can be conveniently expressed as a function of the standard deviation of the signal of interest in the blank and the coefficients of the fitted model $a$ and $b$. Moreover, since coefficient $c$ in equation 5 reflects the background signal when the drug signal is absent, the standard error of $c$ derived from the nonlinear model fit can be used to approximate $\sigma_{Blank}$.

As in the case of the linear model, the nonlinear regression was weighted by the inverse of the variance of each calibration point to limit the impact of the signal heteroscedasticity on the fitted model.

##### 1.3. MALDI-TOF and –FTICR instrument settings

MALDI-TOF ultrafleXtreme instrument parameters: 500 laser shots per position; mass range m/z 300-2000; detector gain set to 50 V under the voltage recommended by detector check.

MALDI-FTICR SolariX instrument parameters: using absorption mode, a 1M data point transient, pixel sizes as for MALDI-TOF, but 10 laser shots per pixel. Data acquisition parameters in ftmsControl (Bruker Daltonics): Source Optics and Ion transfer (Plate offset 100C, Deflector Plate 200 V, Funnel 1 150 V, Skimmer 1 15 V, Funnel RF Amplitude 70 Vpp); Octopole (Frequency 5 MHz, RF Amplitude 350 Vpp); Collision Cell (RF Frequency 2 MHz, RF Amplitude 1000 Vpp); Transfer Optics (Time of Flight 1.0 ms, Frequency 4 MHz, FR Amplitude 300 Vpp); Quadrupole (Q1 Mass 350 m/z); Excitation Mode (Sweep Excitation, Sweep Step time 15 µs).

##### 1.4. Tissue Extraction and Imatinib Quantification by UPHPLC-MS Analysis

Four 8 µm-cryosections of patient samples were collected and stored at -80 °C until analysis. For tissue extraction, 500 µL of 50% MeOH/$H_2O$ + 50 nM imatinib-D8 (D8 solvent) were added. Tubes were vortexed for 30 s and incubated for 10 min. in an ultrasonic bath filled with ice water. Samples were then centrifuged at 10,000 x g for 5 min at 4 °C). 200 µL of this extract were mixed with 800 µL D8 solvent, and 500 µL of diluted extract were transferred to a MiniUniPrep filter vial (pore size 0.2 µm; Agilent, Santa Clara, USA) and placed in a cooled autosampler (4 °C). Imatinib content in patient samples was analyzed on a 1290 Infinity UPLC (Agilent) coupled to an Impact II QToF MS with electrospray ionization (Bruker Daltonics). The HPLC-MS system was controlled by HyStar 3.2 software (Bruker Daltonics). Chromatography was performed on a Zorbax

Eclipse Plus C18 Rapid Resolution HD 2.1 x 50 mm 1.8 µm column at 40 °C, guarded by a Zorbax Eclipse XDB-C18 2.1 x 5 mm 1.8 µm column (Agilent). The flow rate was set to 500 µL/min, and the column was equilibrated with MeOH:water (60:40, v/v) + 0.1% FA. A sample volume of 10 µL was injected and eluted with a gradient between solvent A (MeOH + 0.1% FA), and solvent B (Water + 0.1% FA) as follows:

| Time [min] | A [%] | B [%] | Flow [mL/min] |
|---|---|---|---|
| 0.00 | 40 | 60 | 0.500 |
| 6.00 | 100 | 0 | 0.500 |
| 8.00 | 100 | 0 | 0.500 |
| 10.00 | 40 | 60 | 0.500 |

The following conditions were used for ionization and ion transfer into the mass analyzer: Source conditions: End plate offset 450 V; capillary voltage 4.5 kV; source temperature 220 °C; dry gas flow 8.0 L/min; nebulizer pressure 1.8 bar. Transfer conditions: Funnel 1 RF 150 V; Funnel 2 RF 200 V; Hexapole RF 50 V; Quadrupole ion energy 4 eV; Collision energy 7 eV; Collision RF 800 eV; Transfer time 30 µs; Pre pulse storage 10 µs. Imatinib was detected and quantified as doubly-protonated adduct $[M]^{2+}$ at m/z 247.634. For quantification, all imatinib signals were first normalized to imatinib-D8, added as IS in the extraction step and afterwards quantified over the mean of external standard curves measured before, in the middle of and at the end of every replicate series. For external calibration curves, a 100 µM stock of imatinib in D8 solvent was serially diluted in 10-fold steps in D8 solvent to a lowest concentration of 0.01 µM. 10 µL of 1 µM, 0.1 µM and 0.01 µM imatinib in D8 solvent were injected, with said solvent serving as blank.

### 2. Supplemental Tables

**Supplemental Table 1:** Clinical and pathological data of the patient series. Independent of therapeutic dosing, all patients received daily dosage of 400-800 mg imatinib until the day of surgery (n.i. = not identified; HPF = high power field).

| Organ (ID) | Metastasis (m) or primary tumor (p) | Sex | Age at surgery | Imatinib regime (months at that dose) (mg/day) | Mutations | | mitosis activity |
|---|---|---|---|---|---|---|---|
| | | | | | C-Kit | PDGFRA | |
| Liver (A) | m | female | 44 | 91<br>400 | Ex9;11;13;17 | Ex18 | 50/50 HPF |
| Liver (C) | m | male | 56 | 15/11/62<br>400/600/800 | Ex17 | --- | n.i. |
| Liver (G) | m | female | 47 | 17<br>800 | Ex9;11;13;17 | Ex18 | n.i. |
| Liver (J) | m | male | 50 | 82<br>800 | Ex11;17 | Ex18 | 36/50 HPF |
| Liver (K) | m | male | 45 | 45<br>400 | Ex11;13 | --- | 11/10 HPF |
| Liver (L) | m | female | 49 | 7/15<br>400/800 | Ex11 | --- | 1/50 HPF < 7/1 HPF |
| Liver (M) | m | male | 56 | 33/25<br>400/600 | n.i. | n.i. | 38/50 HPF |
| Liver (U) | m | male | 62 | n.i.<br>n.i. | Ex11 | --- | 3/50 HPF |
| Liver (V) | m | female | 67 | 31<br>400 | Ex9;11;13;17 | Ex12;14;18 | 92/50 HPF |
| Liver (W) | m | male | 73 | 89<br>400 | Ex9;11;13;17 | Ex18 | 33/50 HPF |
| Liver (W) | m | male | 73 | 88<br>400 | --- | --- | 33/50 HPF |
| Liver (Z) | m | male | 49 | n.i<br>n.i. | Ex11;13 | Ex18 | 65/50 HPF |
| Peritoneum (I) | m | female | 42 | 11<br>400 | Ex11 | --- | 12/50 HPF |
| Peritoneum (N) | m | male | 84 | 72<br>400 | Ex13 | --- | 85/50 HPF |
| Peritoneum (P) | m | male | 62 | 62<br>400 | Ex12 | --- | 4/50 HPF |
| Peritoneum (Q) | m | male | 45 | 8<br>400 | Ex11 | --- | 147/50 HPF |
| Peritoneum (R) | m | male | 76 | 36<br>400 | Ex9;11;13;17 | Ex18 | 30/50 HPF |
| Stomach (D) | p | female | 76 | 10<br>400 | Ex11 | --- | Not vital |
| Stomach (F) | p | male | 52 | 6<br>400 | Ex11 | --- | <1/50 HPF |
| Stomach (T) | p | male | 69 | 13<br>n.i. | --- | --- | 90/50 HPF |
| Stomach (X) | p | female | 87 | 7<br>400 | Ex9;11;13;17 | Ex18 | 10/50 HPF |
| Thorax wall (E) | m | male | 76 | 26/5<br>400/800 | Ex11;117 | Ex12;14;18 | 115/50 HPF |
| Abdomen (Y) | p | male | 43 | n.i.<br>400 | --- | Ex15 | 10/50 HPF |
| Intestine (B) | p | male | 66 | 10<br>400 | Ex13 | --- | 5/50 HPF |
| Intestine (AA) | p | male | 44 | 10<br>800 | Ex9 | --- | 4/50 HPF |
| Intestine (AB) | m | male | 63 | 50<br>400 | Ex9;11;117 | Ex12;14;18 | 17/50 HPF |
| Lung (S) | m | male | 74 | 28<br>400 | Ex11 | --- | 40/50 HPF |
| Colon (O) | p | female | 80 | 12<br>400 | Ex9;11;13;17 | --- | >5/50 HPF |

#### 3. Supplemental Figures

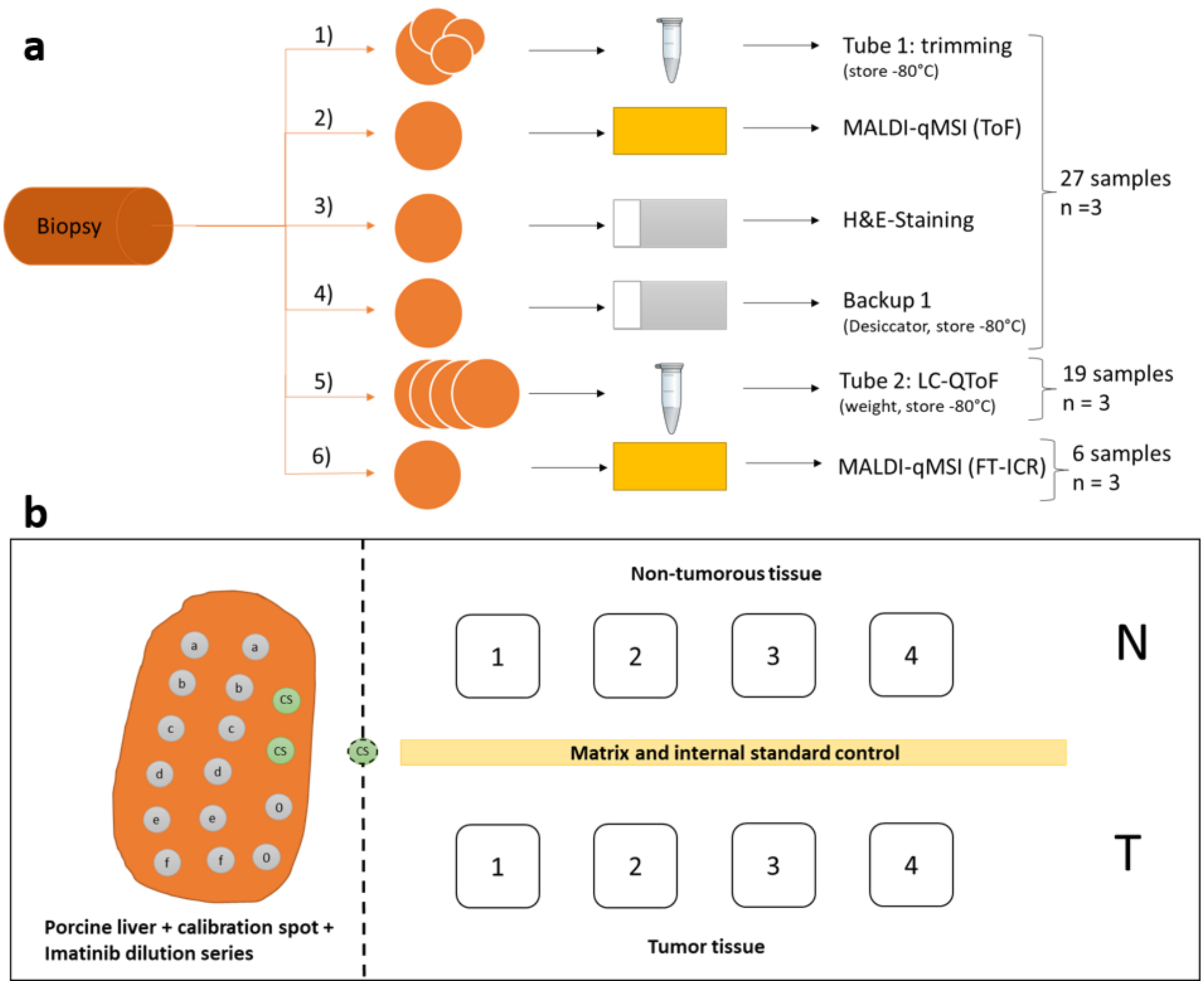


**Supplemental Figure 1:** Experimental design for clinical quantitative MS imaging (qMSI) of imatinib in gastrointestinal tumor (GIST) cohort. **a)** Cutting scheme for the samples**.** After trimming (step 1), the first 8-µm slice of each sample was mounted onto a gold target for MALDI-TOF-qMSI (step 2). Two consecutive slices were then put onto glass slides for standard histological analysis (steps 3/4), followed by the collection of four slices for quantitative drug determination by UPLC-ESI-QTOF-MS (step 5). Steps 1-5 were done for all samples in triplicate. At last, some exemplary slices were mounted on conductive ITO-slides for additional MALDI-qMSI with a high-resolving FTICR (step 6). **b)** Layout of the sample slide used for MALDI-qMSI**.** Duplicate spots per dilution of imatinib (a = 25 pmol; b = 12.5 pmol; c = 6.25 pmol; d = 3.125 pmol; e = 1.5625 pmol; f = 0.78125 pmol; 0 = vehicle control) as well as two spots for mass calibration (CS; green) were spotted onto a porcine liver section mounted on the left side of each slice. Per slide, normal tissue (N; upper row) and tumor tissue (T; lower row) of four patients were analyzed. Between normal and tumor samples, a lane of matrix + internal standard D8-imatinib was measured for calculation of tissue extinction coefficients (TEC).

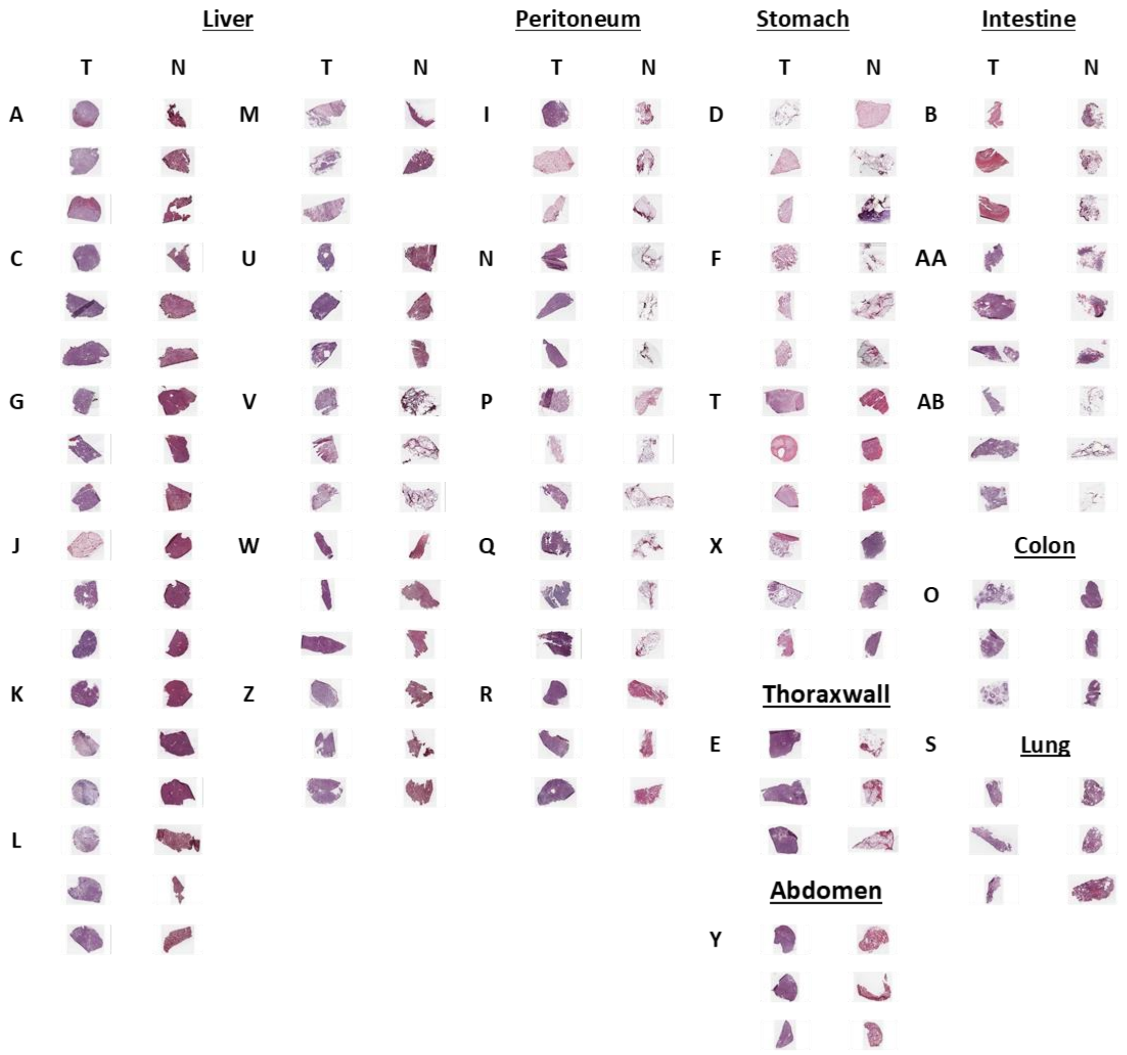


**Supplemental Figure 2:** H&E images of all tumor (T) and normal (N) samples from GIST patients**.** Three cryosections were prepared per tissue sample/patient, and patients are coded by single or double letters (see **Supplemental Table 1**).

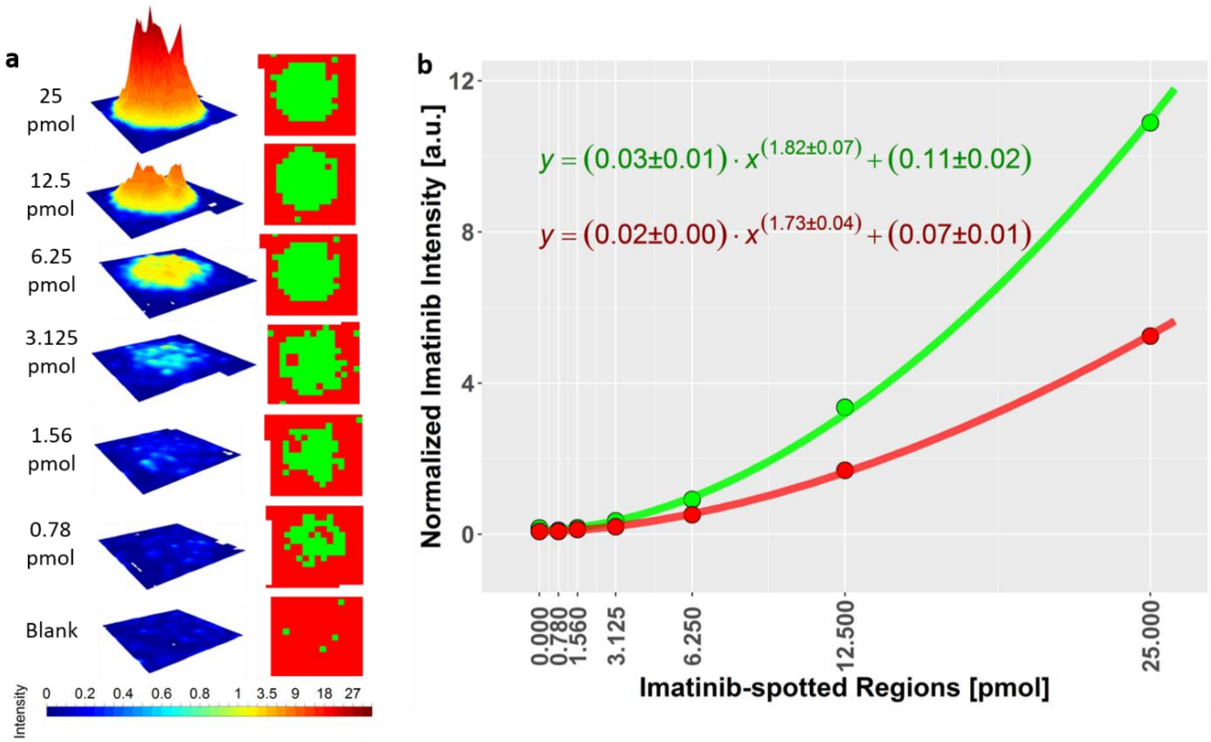


**Supplemental Figure 3:** Imatinib dilution series for one of the datasets shown as **a**) surface- and peak detection plots. **b**) Fitted nonlinear calibration curves based on drug-bearing pixels (green curve) and user-defined areas (red curve) for a chosen MALDI-TOF-qMSI dilution series. Solid circles represent the mean drug intensity of pixels depending on the method used.

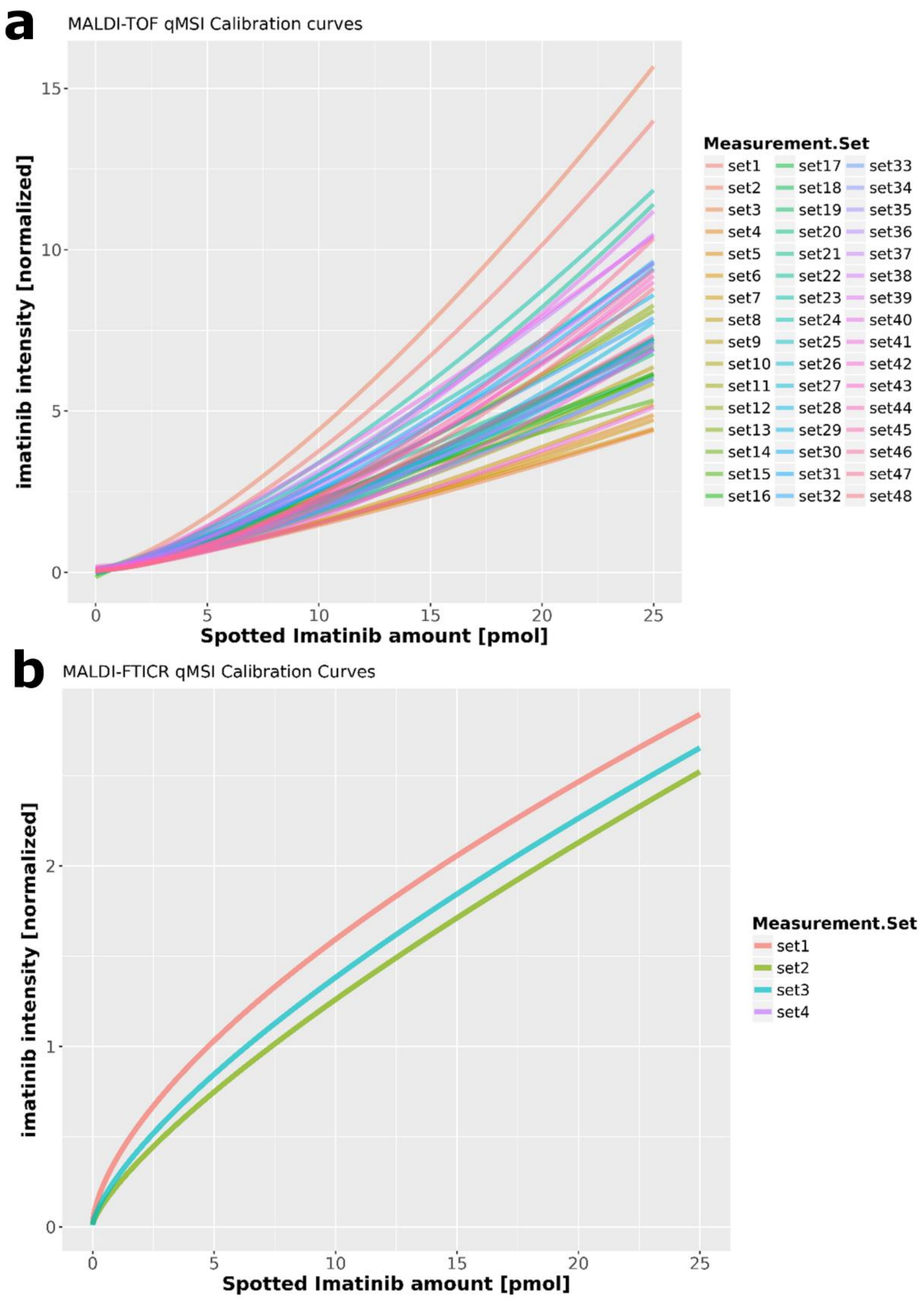


**Supplemental Figure 4:** Ilustration of the nonlinearity response observed in all (**a**) 48 and (**b**) 3 calibration curves obtained for MALDI-TOF-qMSI and MALDI-FTICR-qMSI measurement sets, respectively.

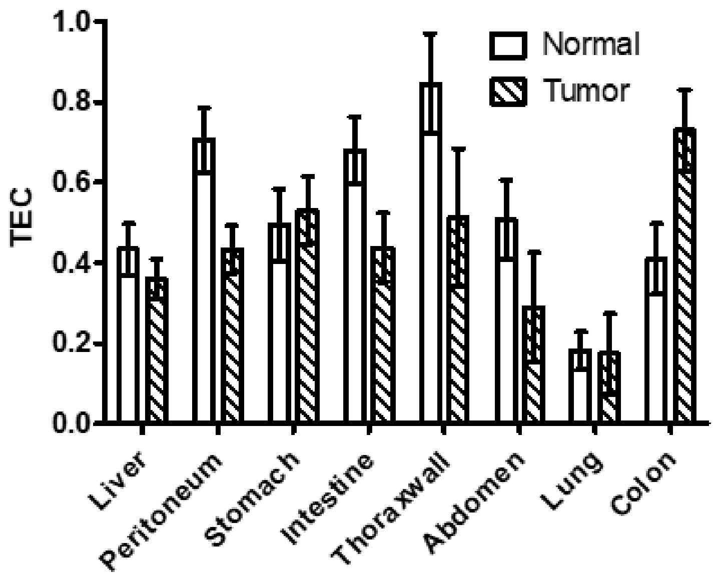


**Supplemental Figure 5:** Tissue-averaged Tissue Extinction coefficient (TEC) calculated across all tissue types based on MALDI-TOF-qMSI.

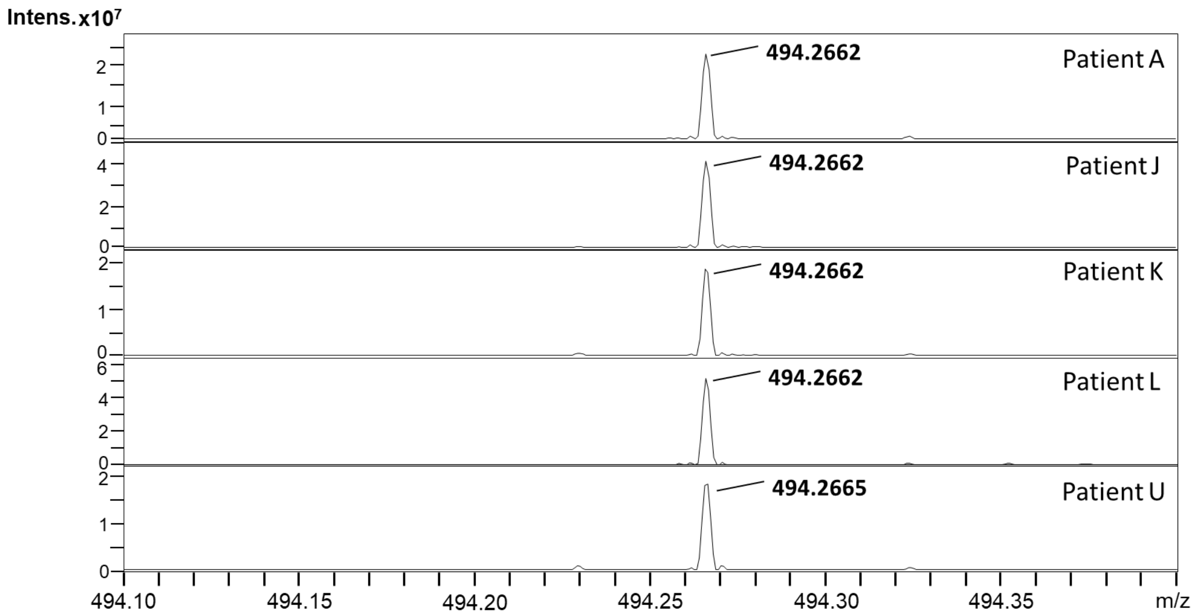


**Supplemental Figure 6:** MALDI-FTICR high mass resolution spectra to exclude molecular interference. No other molecules which could impact the lower resolution MALDI-ToF-qMSI data were observed within $100\ ppm$. The mass resolution of the detected Imatinib peak above evaluates to $185000$.

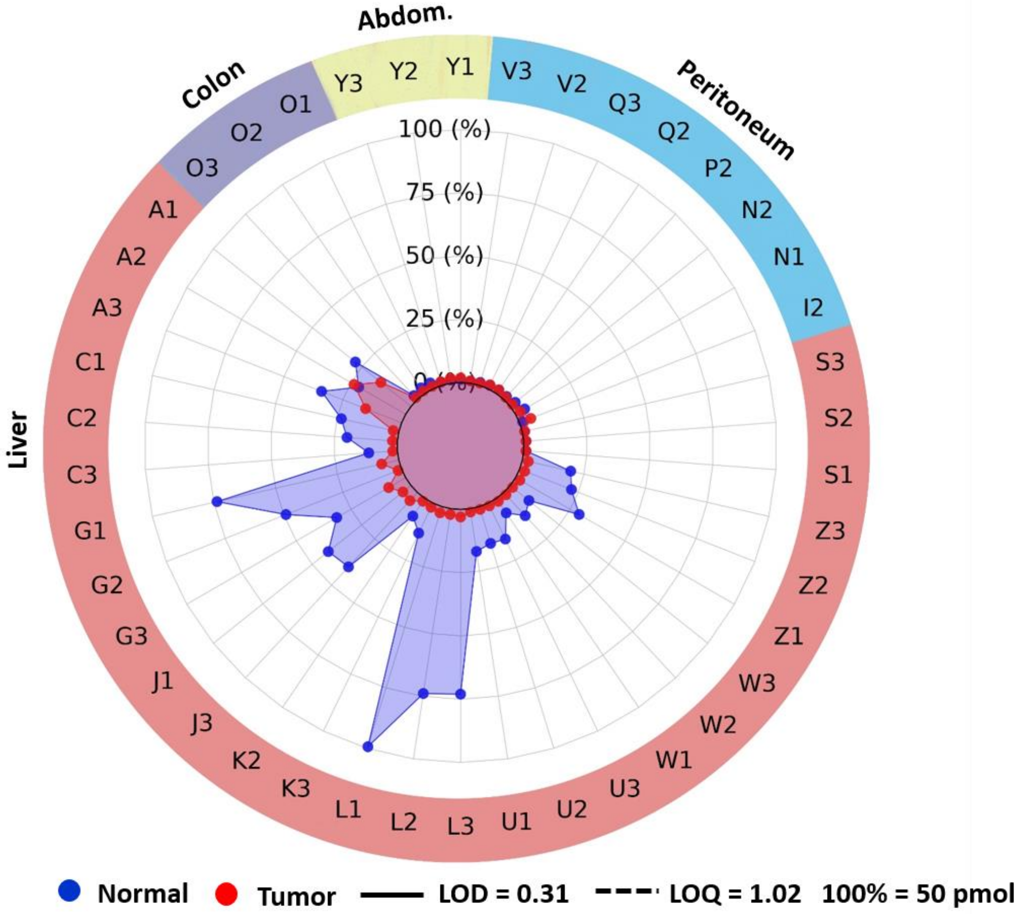


**Supplemental Figure 7:** Imatinib content in $pmol/section$ as quantified by UPLC-ESI-QTOF-MS for 18 patient-derived GIST ("Tumor"; red) and surrounding non-tumor ("Normal"; blue) tissue sections (N = 3 when vaialble; LOD = 0.31 pmol/section; LOQ = 1.02 pmol/secion).
